# Virological characteristics of the SARS-CoV-2 Omicron XBB.1.16 variant

**DOI:** 10.1101/2023.04.06.535883

**Authors:** Daichi Yamasoba, Keiya Uriu, Arnon Plianchaisuk, Yusuke Kosugi, Lin Pan, Jiri Zahradnik, The Genotype to Phenotype Japan (G2P-Japan) Consortium, Jumpei Ito, Kei Sato

## Abstract

At the end of March 2023, XBB.1.16, a SARS-CoV-2 omicron XBB subvariant, emerged and was detected in various countries. Compared to XBB.1.5, XBB.1.16 has two substitutions in the S protein: E180V is in the N-terminal domain, and T478R in the receptor-binding domain (RBD). We first show that XBB.1.16 had an effective reproductive number (Re) that was 1.27- and 1.17-fold higher than the parental XBB.1 and XBB.1.5, respectively, suggesting that XBB.1.16 will spread worldwide in the near future. In fact, the WHO classified XBB.1.16 as a variant under monitoring on March 30, 2023. Neutralization assays demonstrated the robust resistance of XBB.1.16 to breakthrough infection sera of BA.2 (18-fold versus B.1.1) and BA.5 (37-fold versus B.1.1). We then used six clinically-available monoclonal antibodies and showed that only sotrovimab exhibits antiviral activity against XBB subvariants, including XBB.1.16. Our results suggest that, similar to XBB.1 and XBB.1.5, XBB.1.16 is robustly resistant to a variety of anti-SARS-CoV-2 antibodies. Our multiscale investigations suggest that XBB.1.16 that XBB.1.16 has a greater growth advantage in the human population compared to XBB.1 and XBB.1.5, while the ability of XBB.1.16 to exhibit profound immune evasion is comparable to XBB.1 and XBB.1.5. The increased fitness of XBB.1.16 may be due to (1) different antigenicity than XBB.1.5; and/or (2) the mutations in the non-S viral protein(s) that may contribute to increased viral growth efficiency.

## Text

In late February 2023, certain sublineages of the SARS-CoV-2 omicron XBB variant harboring the F486P substitution in the spike (S) protein (e.g. XBB.1.5 and XBB.1.9) predominated worldwide (https://nextstrain.org/ncov/gisaid/global/6m). Subsequently, XBB.1.16, an XBB sublineage, emerged and was detected in various countries. Compared to XBB.1.5, XBB.1.16 has two substitutions in the S protein: E180V in the N-terminal domain, and T478R in the receptor-binding domain (RBD) (**Figure S1A**). XBB.1.16 outcompeted other variants in India by the end of March 2023 (**Figure S1B**). Notably, XBB.1.16 had an effective reproductive number (R_e_) that was 1.27- and 1.17-fold higher than the parental XBB.1 and XBB.1.5, respectively, suggesting that XBB.1.16 will spread worldwide in the near future (**Figure S1C**). In fact, the WHO classified XBB.1.16 as a variant under monitoring on March 30, 2023.^1^

We next investigated the virological features of XBB.1.16. Yeast surface display assay^2^ showed the dissociation constant (K_D_) of XBB.1.16 RBD to the human ACE2 receptor is significantly (2.4-fold) higher than that of XBB.1.5 RBD, while the K_D_ of XBB.1.16 RBD is significantly (1.8-fold) lower than that of XBB.1 RBD (**Figure S1D**). These results suggest the binding affinity of XBB.1.16 RBD to ACE2 is higher than that of XBB.1 RBD and lower than that of XBB.1.5 RBD. Pseudovirus experiments showed higher infectivity of XBB.1.5 compared to the parental XBB.1, which is consistent with our previous study (**Figure S1E**).^3^ In contrast, the pseudovirus infectivity of XBB.1.16 was comparable to that of XBB.1 (**Figure S1E**). The S:T478R substitution significantly increased infectivity, while the S:E180V substitution significantly decreased infectivity (**Figure S1E**). The acquisition of two combination mutations in the S protein, one that evades antiviral immunity and attenuates infectivity (e.g., F486V, G446S, Y144del), and another that increases infectivity (e.g., L452R, N460K, V83A) is a strategy of Omicron evolution previously observed in BA.5^4^, BA.2.75^5^, and XBB.1.^6^ Our findings suggest that XBB.1.16 possibly follows the evolutionary pattern of previous Omicron variants.

Neutralization assays demonstrated the robust resistance of XBB.1.16 to breakthrough infection sera of BA.2 (18-fold versus B.1.1; **Figure S1F**) and BA.5 (37-fold versus B.1.1; **Figure S1G**). On the other hand, the sensitivity of XBB.1.16 to convalescent sera of XBB.1-infected hamsters^6^ was comparable to those of XBB.1 and XBB.1.5 (**Figure S1H**). We then used six clinically-available monoclonal antibodies and showed that only sotrovimab exhibits antiviral activity against XBB subvariants, including XBB.1.16 (**Table S1**). Our results suggest that XBB.1 and XBB.1.5, XBB.1.16 is robustly resistant to a variety of anti-SARS-CoV-2 antibodies. Finally, antigenic cartography based on our results (**Figures S1F-H**) showed that the antigenicity of XBB.1.16 is different from that of XBB.1.5, and rather relatively close to that of XBB.1 (**Figure S1I**).

Altogether, our data suggest that XBB.1.16 has a greater growth advantage in the human population compared to XBB.1 and XBB.1.5, while the ability of XBB.1.16 to exhibit profound immune evasion is comparable to XBB.1 and XBB.1.5. The increased fitness of XBB.1.16 may be due to (1) different antigenicity from XBB.1.5; and/or (2) the mutations in the non-S viral protein(s) that may contribute to increased viral growth efficiency.

## Grants

Supported in part by AMED SCARDA Japan Initiative for World-leading Vaccine Research and Development Centers “UTOPIA” (JP223fa627001, to Kei Sato), AMED SCARDA Program on R&D of new generation vaccine including new modality application (JP223fa727002, to Kei Sato); AMED Research Program on Emerging and Re-emerging Infectious Diseases (JP22fk0108146, to Kei Sato; JP21fk0108494 to G2P-Japan Consortium and Kei Sato; JP21fk0108425, to Kei Sato; JP21fk0108432, to Kei Sato; JP22fk0108511, to G2P-Japan Consortium and Kei Sato; JP22fk0108516, to Kei Sato; JP22fk0108506, to Kei Sato); AMED Research Program on HIV/AIDS (JP22fk0410039, to Kei Sato); JST PRESTO (JPMJPR22R1, to Jumpei Ito); JST CREST (JPMJCR20H4, to Kei Sato); JSPS KAKENHI Grant-in-Aid for Early-Career Scientists (20K15767, to Jumpei Ito; 23K14526, to Jumpei Ito); JSPS Core-to-Core Program (A. Advanced Research Networks) (JPJSCCA20190008, Kei Sato); JSPS Research Fellow DC2 (22J11578, to Keiya Uriu); The Tokyo Biochemical Research Foundation (to Kei Sato); and the project of National Institute of Virology and Bacteriology, Programme EXCELES, funded by the European Union, Next Generation EU (LX22NPO5103, to Jiri Zahradnik).

## Declaration of interest

We declare no competing interests.

## Notes

Conflict of interest: The authors declare that no competing interests exist.

### Competing Interest Statement

The authors have declared no competing interest.

### Summary of Updates

Table S1 is revised - some parameters were replaced with correct ones.

## References

1. WHO. “Tracking SARS-CoV-2 variants (March 30, 2023)” https://www.who.int/en/activities/tracking-SARS-CoV-2-variants. 2022.

2. Zahradnik J, Marciano S, Shemesh M, et al. SARS-CoV-2 variant prediction and antiviral drug design are enabled by RBD in vitro evolution. Nat Microbiol 2021; 6(9): 1188–98.

3. Uriu K, Ito J, Zahradnik J, et al. Enhanced transmissibility, infectivity, and immune resistance of the SARS-CoV-2 omicron XBB.1.5 variant. Lancet Infect Dis 2023; 23(3): 280–1.

4. Kimura I, Yamasoba D, Tamura T, et al. Virological characteristics of the novel SARS-CoV-2 Omicron variants including BA.4 and BA.5. Cell 2022; 185(21): 3992–4007.e16.

5. Saito A, Tamura T, Zahradnik J, et al. Virological characteristics of the SARS-CoV-2 Omicron BA.2.75 variant. Cell Host Microbe 2022; 30(11): 1540–55.e15.

6. Tamura T, Ito J, Uriu K, et al. Virological characteristics of the SARS-CoV-2 XBB variant derived from recombination of two Omicron subvariants. BioRxiv 2022: doi: https://doi.org/10.1101/2022.12.27.521986.

