## Appendix for "Virological characteristics of the SARS-CoV-2 Omicron XBB.1.16 variant"

**Supplementary Appendix**

**Table of Contents**

**Contents** Page


| | |
|---|---|
| **Materials and Methods** | 2-5 |
| Ethics statement | |
| Human serum collection | |
| Epidemic dynamic analysis | |
| Plasmid construction | |
| Yeast surface display | |
| Cell culture | |
| Pseudovirus infection | |
| Neutralization assay | |
| Data availability | |
| **Table S1.** IC50s of six therapeutic monoclonal antibodies against XBB.1.16 | 6 |
| **Table S2.** Human sera used in this study | 7 |
| **Table S3.** Estimated relative $R_e$ and epidemic dynamics modeling parameters of the representative SARS-CoV-2 Omicron sublineages spreading in India from October 1, 2022 to March 5, 2023. | 8 |
| **Table S4.** Primers used in this study | 9 |
| **Figure S1.** Virological features of XBB.1.16 | 10-11 |
| **Consortia** | 12 |
| **Acknowledgments** | 13 |
| **Supplemental References** | 14-15 |

### Materials and Methods

#### Ethics statement

All protocols involving specimens from human subjects recruited at Interpark Kuramochi Clinic was reviewed and approved by the Institutional Review Board of Interpark Kuramochi Clinic (approval ID: G2021-004). All human subjects provided written informed consent. All protocols for the use of human specimens were reviewed and approved by the Institutional Review Boards of The Institute of Medical Science, The University of Tokyo (approval IDs: 2021-1-0416 and 2021-18-0617).

#### Human serum collection

Convalescent sera were collected from fully vaccinated individuals who had been infected with BA.2 (9 2-dose vaccinated and 4 3-dose vaccinated; time interval between the last vaccination and infection, 4–299 days； 11–61 days after testing. n=13 in total; average age: 45 years, range: 24–82 years, 62% male) (**Figure S1F**), and fully vaccinated individuals who had been infected with BA.5 (1 2-dose vaccinated, 13 3-dose vaccinated and 1 4-dose vaccinated; time interval between the last vaccination and infection, 66–310 days; 10–23 days after testing. n=15 in total; average age: 55 years, range: 25–73 years, 47% male) (**Figure S1G**). The SARS-CoV-2 variants were identified as previously described.[1-3] Sera were inactivated at 56°C for 30 minutes and stored at –80°C until use. The details of the convalescent sera are summarized in **Table S2**.

#### Method

#### Epidemic dynamics analysis and mutation frequency calculation

We modeled the epidemic dynamics of SARS-CoV-2 lineage spreading in India from October 1, 2022 to March 5, 2023 with the viral genomic surveillance data deposited in the GISAID database (https://www.gisaid.org/; downloaded on March 25, 2023). We excluded the data of the SARS-CoV-2 isolate that i) lacks collection date and PANGO lineage information; ii) was retrieved from non-human animals; iii) was sampled by quarantine; iv) was sampled from the original passage; and v) whose genomic sequence is not longer than 28,000 base pairs and contains >2% of unknown (N) nucleotides. We defined XBB.1 harboring the S:E180V+G252V+T478R+F486P substitutions as XBB.1.16. Only the SARS-CoV-2 lineages with >50 genomic sequences were kept for epidemic dynamics analysis (**Figures S1B and S1C**). SARS-CoV-2 in XBB sublineages that have spread in India during the specified time include those in XBB, XBB.1, XBB.1.5, XBB.1.16, XBB.2, and XBB.3 sublineages. We treated SARS-CoV-2 in XBB.1, XBB.1.5, XBB.2, and XBB.3 sublineages lacking their signature S:G252V, S:G252V+F486P, S:D253G, and NSP1:G82D+NSP14:I474V substitutions, respectively, as misannotated and subsequently discarded them. We also treated SARS-CoV-2 in XBB sublineage with S:F486P as misannotated and discarded them. Epidemic dynamics and a relative $R_e$ of each viral lineage were estimated based on the Bayesian multinomial logistic model, described in our previous study.[2] Briefly, we used the model to estimate the logistic slope parameter $\beta_l$ for each lineage and then calculated a relative $R_e$ for each lineage ($r_l$) as $r_l = exp(\gamma\beta_l)$ where $\gamma$ is the average viral generation time (2.1 days) (http://sonorouschocolate.com/covid19/index.php?title=Estimating_Generation_Time_Of_Omicron). For parameter estimation, the intercept and slope parameters of XBB.1 were fixed at 0. The relative $R_e$ of XBB.1 was fixed at 1, and that of other lineage was estimated with respect to that of XBB.1. Parameter estimation was performed by using the Markov chain Monte Carlo (MCMC) approach implemented in CmdStan v2.31.0 (https://mc-stan.org) accessed through the CmdStanr v0.5.3 R interface (https://mc-stan.org/cmdstanr/). Four independent 5,000-step MCMC chains were run including 1,000-step warmup iterations. We confirmed that an estimated $\hat{R}$ convergence diagnostic value is <1.01 and an effective sampling size is >200, indicating that all runs were successfully convergent (**Table S3**). Only the results for XBB.1, XBB.1.5, and XBB.1.16 are shown in **Figures S1B and S1C**, and estimated values for other lineages are summarized in **Table S3**. Mutation frequency of each lineage was calculated by dividing the number of sequences harboring the substitution of interest with the total number of sequences in each lineage. The heatmap of mutation frequencies was created by using ComplexHeatmap v2.14.0.[4]

**Plasmid construction**

Plasmids expressing the SARS-CoV-2 spike proteins of the parental D614G (B.1.1), Omicron BA.2, BA.5, XBB.1, and XBB.1.5 were prepared in our previous studies.[1,2,5-10] Plasmids expressing the spike protein of XBB.1.5 and its derivative were generated by site-directed overlap extension PCR using pC-SARS2-S XBB.1[9] as the template and the primers listed in **Table S4**. The resulting PCR fragment was subcloned into the KpnI-NotI site of the pCAGGS vector[11] using In-Fusion® HD Cloning Kit (Takara, Cat# Z9650N). Nucleotide sequences were determined by DNA sequencing services (Eurofins), and the sequence data were analyzed by Sequencher v5.1 software (Gene Codes Corporation). Plasmids expressing anti-SARS-CoV-2 monoclonal antibodies (bebtelovimab, casirivimab, cilgavimab, imdevimab, sotrovimab, tixagevimab) were prepared as previously described.[7]

**Yeast surface display**

Yeast surface display binding analyses for the spike receptor-binding domains of BA.2, XBB and XBB.1.5 (residues 333–527) were performed as previously described.[1-3,5,8,9,12-14] Cerevisiae EBY100 yeasts and pJYDC1 plasmids with RBD genes cloned between the NdeI and BamHI sites (Addgene, Cat# 162458) as previously described.[12,13] Transformed yeasts were grown in SDCAA media (220 rpm, 30°C) and expressed overnight in expression media supplemented with bilirubin (10 nM DMSO solubilized, Sigma-Aldrich, Cat# 14370) after inoculation to OD600 0.7–1.0 (220 rpm, 20°C).[12,13] Aliquots of expressed yeast cells (100 µl) were washed in ice-cold PBSB buffer (PBS with 1 mg/ml BSA) and incubated in a series of CF®640R succinimidyl ester labeled (Biotium, Cat# 92108) ACE2 peptidase domain (residues 18–740) concentrations, PBSB buffer and 1 nM bilirubin for 8 to 12 hours. After incubation, the unbound fraction was washed by ice-cold PBSB buffer and yeasts were transferred into a 96-well plate (Thermo Fisher Scientific, Cat# 268200) and 30,000 events in gated population were automatically acquired by a CytoFLEX S Flow Cytometer (Beckman Coulter, USA, Cat#. N0-V4-B2-Y4) with FITC channel data for eUnaG2 fluorescence (Abs/Em maxima 498/527 nm) and APC channel for CF640 fluorescence (Abs/Em maxima 642/662 nm) setting. Gating, analysis and fitting with nonlinear least-squares regression using Python v3.7 protocols were described previously.[1-3,5,8,9,12-14]

**Cell culture**

HEK293T cells (a human embryonic kidney cell line; ATCC CRL-3216) and HOS-ACE2/TMPRSS2 cells (kindly provided by Dr. Kenzo Tokunaga),[15,16] a derivative of HOS cells (a human osteosarcoma cell line; ATCC CRL-1543) stably expressing human ACE2 and TMPRSS2, were maintained in Dulbecco’s modified Eagle's medium (DMEM) (high glucose) (Wako, Cat# 044-29765) containing 10% fetal bovine serum (FBS) (Sigma-Aldrich Cat# 172012-500ML), 100 units penicillin and 100 ug/ml streptomycin (PS) (Sigma-Aldrich, Cat# P4333-100ML).

**Pseudovirus infection**

Pseudoviruses were prepared as previously described. Briefly, lentivirus (HIV-1)-based, luciferase-expressing reporter viruses were pseudotyped with the SARS-CoV-2 spikes. HEK293T cells ($1 \times 10^6$ cells) were cotransfected with 1 µg psPAX2-IN/HiBiT, 1 µg pWPI-Luc2, and 500 ng plasmids expressing parental S or its derivatives using PEI Max (Polysciences, Cat# 24765-1) according to the manufacturer's protocol. Two days post transfection, the culture supernatants were harvested and centrifuged. The amount of pseudoviruses prepared was quantified by the HiBiT assay using Nano Glo HiBiT lytic detection system (Promega,Cat# N3040) as previously described. To measure viral infectivity, the same amount of pseudoviruses (normalized to the HiBiT value, which indicates the amount of p24 HIV-1 antigen) was inoculated into HOS-ACE2/TMPRSS2 cells. Two days post infection, the infected cells were lysed with a Bright-Glo luciferase assay system (Promega, Cat# E2620), and the luminescent signal was measured using a GloMax explorer multimode microplate reader 3500 (Promega). The pseudoviruses were stored at –80°C until use.

**Neutralization assay**

Pseudoviruses were prepared as previously described.[1-3,6-9,14,15,17-20] Briefly, lentivirus (HIV-1)-based, luciferase-expressing reporter viruses were pseudotyped with the SARS-CoV-2 spikes. HEK293T cells ($2 \times 10^6$ cells) were cotransfected with 1 µg psPAX2-IN/HiBiT,[21] 1 µg pWPI-Luc2,[21] and 500 ng plasmids expressing parental S or its derivatives using PEI Max (Polysciences,

Cat# 24765-1) according to the manufacturer's protocol. Two days post transfection, the culture supernatants were harvested and centrifuged. The pseudoviruses were stored at –80°C until use. Neutralization assays were performed as previously described.[1-3,6-10,14,15,17-20] Six monoclonal antibodies (bebtelovimab, casirivimab, cilgavimab, imdevimab, sotrovimab, tixagevimab) were prepared as previously described.[7] The SARS-CoV-2 spike pseudoviruses (counting ~20,000 relative light units) were incubated with serially diluted (120-fold to 87,480-fold dilution at the final concentration) heat-inactivated sera or serially diluted monoclonal antibodies at 37°C for 1 hour. Pseudoviruses without sera were or monoclonal antibodies included as controls. Then, an 20 µl mixture of pseudovirus and serum or monoclonal antibody was added to HOS-ACE2/TMPRSS2 cells (10,000 cells/100 µl) in a 96-well white plate. Two days post infection, the infected cells were lysed with a Bright-Glo luciferase assay system (Promega, Cat# E2620), and the luminescent signal was measured using a GloMax explorer multimode microplate reader 3500 (Promega). The assay of each serum or monoclonal antibody sample was performed in triplicate, and the 50% neutralization titer (NT50) or 50% inhibition concentration (IC50) was calculated using Prism 9 (GraphPad Software).

Principal component analysis representing the antigenicity of the S proteins was performed (**Figure S1I**). The NT50 values were scaled, and subsequently, principal component analysis was performed using the PCATools v2.10.0 on R v4.2.2 (https://www.r-project.org/).

**Data availability**

The GISAID dataset used in this study is available from the GISAID database (https://www.gisaid.org; EPI_SET_230403zh). The supplemental table for the EPI_SET_230403zh dataset is available in the GitHub repository (https://github.com/TheSatoLab/XBB.1.16_short).

**Table S1. IC50s of six therapeutic monoclonal antibodies against XBB.1.16.**

| | B.1.1 | BA.2 | BA.5 | XBB.1 |
|---|---|---|---|---|
| Bebtelovimab | 5.2 ± 1.7 | 3.5 ± 1.4 | 3.4 ± 1.6 | > 775 |
| Casirivimab | 12 ± 3.1 | > 5042 | > 5042 | > 5042 |
| Cilgavimab | 33 ± 7.5 | 39 ± 3.5 | 692 ± 220 | > 4200 |
| Imdevimab | 35 ± 17 | > 5000 | > 5000 | > 5000 |
| Sotrovimab | 148 ± 130 | 2950 ± 2301 | 2454 ± 1576 | 607 ± 314 |
| Tixagevimab | 5.8 ± 1.8 | 1130 ± 1647 | > 2600 | > 2600 |
| Ronapreve (casirivimab+imdevimab) | 7.9 ± 1.8 | > 4860 | > 4860 | > 4860 |
| Evusheld (cilgavimab+tixagevimab) | 4.3 ± 0.11 | 35 ± 18 | 433 ± 126 | > 2100 |

| | XBB.1.5 | XBB.1.5+E180V | XBB.1.5+T478R | XBB.1.16 |
|---|---|---|---|---|
| Bebtelovimab | > 775 | > 775 | > 775 | > 775 |
| Casirivimab | > 5042 | > 5042 | > 5042 | > 5042 |
| Cilgavimab | > 4200 | > 4200 | > 4200 | > 4200 |
| Imdevimab | > 5000 | > 5000 | > 5000 | > 5000 |
| Sotrovimab | 740 ± 225 | 836 ± 648 | 922 ± 57 | 780 ± 764 |
| Tixagevimab | > 2600 | > 2600 | > 2600 | > 2600 |
| Ronapreve (casirivimab+imdevimab) | > 4860 | > 4860 | > 4860 | > 4860 |
| Evusheld (cilgavimab+tixagevimab) | > 2100 | > 2100 | > 2100 | > 2100 |

Neutralization assay was performed using pseudoviruses harboring the SARS-CoV-2 spike proteins of BA.2, BA.5, XBB.1, XBB.1.5, XBB.1.5+E180V, XBB.1.5+T478R and XBB.1.16 (XBB.1.5 spike: E180V/T478R) or the D614G-harboring B.1.1 lineage virus (B.1.1). Six therapeutic monoclonal antibodies (bebtelovimab, casirivimab, cilgavimab, imdevimab, sotrovimab and tixagevimab) and two antibody cocktails [Ronapreve (casirivimab+imdevimab), Evusheld (cilgavimab+tixagevimab)] were tested. The assay of each antibody was performed in triplicate at each concentration to determine the 50% inhibitory concentration (IC50; ng/ml), and the assay was independently repeated three times. The presented data are expressed as the average ± 95% confidential interval.

**Table S2. Human sera used in this study**

| SARS-CoV-2 infected | Donor ID | Sex | Age | Date of test (YYYY/MM/DD) | Date of sampling (YYYY/MM/DD) | Prior infection? | Prior vaccination? | Vaccine | Date of 1st vaccination (YYYY/MM/DD) | Date of 2nd vaccination (YYYY/MM/DD) | Date of 3rd vaccination (YYYY/MM/DD) | Date of 4rd vaccination (YYYY/MM/DD) |
|---|---|---|---|---|---|---|---|---|---|---|---|---|
| BA.2 | P378 | Male | 43 | 2022/03/28 | 2022/04/10 | No | Yes | BNT162b2 | 2021/10/10 | 2021/10/31 | | |
| BA.2 | P398 | Male | 48 | 2022/04/13 | 2022/04/30 | No | Yes | BNT162b2 | 2021/09/18 | 2021/10/09 | 2022/04/09 | |
| BA.2 | P407 | Male | 29 | 2022/05/01 | 2022/05/12 | No | Yes | mRNA-1273 | 2021/09/13 | 2021/10/11 | | |
| BA.2 | P401 | Male | 35 | 2022/04/22 | 2022/05/05 | No | Yes | BNT162b2 | 2021/09/09 | 2021/09/30 | | |
| BA.2 | P412 | Female | 82 | 2022/05/04 | 2022/05/26 | No | Yes | BNT162b2 | 2021/06/11 | 2021/07/09 | | |
| BA.2 | 6449 | Male | 43 | 2022/04/03 | 2022/04/23 | No | Yes | BNT162b2 | 2021/08/13 | 2021/09/11 | | |
| BA.2 | 6355 | Male | 50 | 2022/04/02 | 2022/04/20 | No | Yes | NA | 2021/04/28 | 2021/05/19 | 2022/01/19 | |
| BA.2 | 6547 | Male | 54 | 2022/04/06 | 2022/04/22 | No | Yes | BNT162b2 | 2021/08/25 | 2021/09/15 | | |
| BA.2 | 7951 | Female | 71 | 2022/04/25 | 2022/05/12 | No | Yes | Mix | 2021/06/20 (BNT162b2) | 2021/07/16 (BNT162b2) | 2022/02/16 (mRNA-1273） | |
| BA.2 | 8645 | Female | 41 | 2022/05/07 | 2022/05/20 | No | Yes | BNT162b2 | 2021/05/23 | 2021/06/13 | 2022/01/20 | |
| BA.2 | 8682 | Female | 25 | 2022/05/08 | 2022/05/24 | No | Yes | BNT162b2 | 2021/09/03 | 2021/09/27 | | |
| BA.2 | 5949 | Male | 24 | 2022/03/22 | 2022/05/22 | No | Yes | mRNA-1273 | 2021/08/05 | 2021/09/02 | | |
| BA.2 | 8796 | Female | 34 | 2022/05/10 | 2022/06/05 | No | Yes | BNT162b2 | 2021/09/26 | 2021/10/17 | | |
| BA.5 | P427 | Female | 49 | 2022/07/06 | 2022/07/25 | No | Yes | NA | 2021/07/30 | 2021/08/25 | 2022/03/18 | |
| BA.5 | P439 | Female | 73 | 2022/07/23 | 2022/08/08 | No | Yes | Mix | 2021/6/19 (BNT162b2) | 2021/7/20 (BNT162b2) | 2022/02/04 (mRNA-1273) | |
| BA.5 | P451 | Female | 55 | 2022/07/29 | 2022/8/12 | No | Yes | BNT162b2 | 2021/04/26 | 2021/05/20 | 2022/01/18 | |
| BA.5 | P456 | Male | 44 | 2022/08/04 | 2022/08/14 | No | Yes | Mix | 2021/8/11 (BNT162b2) | 2021/9/1 (BNT162b2) | 2022/03/13 (mRNA-1273) | |
| BA.5 | P464 | Male | 63 | 2022/08/08 | 2022/08/19 | No | Yes | Mix | 2021/8/8 (BNT162b2) | 2021/8/29 (BNT162b2) | 2022/04/07 (mRNA-1273) | |
| BA.5 | 9341 | Male | 56 | 2022/06/12 | 2022/06/30 | No | Yes | BNT162b2 | 2021/08/10 | 2021/08/31 | 2022/03/18 | |
| BA.5 | 9584 | Male | 55 | 2022/07/08 | 2022/07/25 | No | Yes | BNT162b2 | 2021/07/14 | 2021/08/05 | 2022/03/22 | |
| BA.5 | 11318 | Female | 51 | 2022/07/24 | 2022/08/05 | No | Yes | Mix | 2021/9/1 (BNT162b2) | 2021/9/22 (BNT162b2) | 2022/05/19 (mRNA-1273) | |
| BA.5 | 23S-08 | Male | 25 | 2022/07/23 | 2022/08/08 | No | Yes | BNT162b2 | 2021/04/27 | 2021/05/18 | 2022/01/11 | |
| BA.5 | 10826 | Male | 63 | 2022/07/21 | 2022/08/11 | No | Yes | Mix | 2021/7/27 (BNT162b2) | 2021/8/17 (BNT162b2) | 2022/3/4 (BNT162b2) | 2022/8/9 (mRNA-1273) |
| BA.5 | 11079 | Female | 65 | 2022/07/23 | 2022/08/11 | No | Yes | mRNA-1273 | 2021/07/08 | 2021/08/05 | 2022/03/17 | |
| BA.5 | 14847 | Female | 70 | 2022/08/13 | 2022/08/25 | No | Yes | Mix | 2021/7/13 (BNT162b2) | 2021/8/20 (BNT162b2) | 2022/03/08 (mRNA-1273) | |
| BA.5 | 13180 | Female | 63 | 2022/08/04 | 2022/08/25 | No | Yes | Mix | 2021/7/16 (BNT162b2) | 2021/8/6 (BNT162b2) | 2022/03/08 (mRNA-1273) | |
| BA.5 | 12912 | Male | 64 | 2022/08/02 | 2022/08/25 | No | Yes | mRNA-1273 | 2021/09/02 | 2021/09/30 | 2021/04/01 | |
| BA.5 | 14956 | Female | 33 | 2022/08/13 | 2022/08/28 | No | Yes | BNT162b2 | 2021/09/06 | 2021/10/07 | | |

NA, not applicable.

**Table S3. Estimated relative Re and epidemic dynamics modeling parameters of the representative SARS-CoV-2 Omicron sublineages spreading in India from October 1, 2022 to March 5, 2023**

| PANGO lineage | Relative $R_e$ (posterior values) | | | $\hat{R}$ | Bulk effective sample size | Tail effective sample size |
|---|---|---|---|---|---|---|
| | Mean | $2.5^{th}$ percentile | $97.5^{th}$ percentile | | | |
| XBB.1.16 | 1.266 | 1.211 | 1.332 | 1.000 | 13464.5 | 9909.5 |
| XBB.1.5 | 1.083 | 1.069 | 1.099 | 1.000 | 11788.5 | 10606.8 |
| BA.2.10.1 | 1.029 | 1.019 | 1.039 | 1.000 | 9911.2 | 11336.9 |
| XBB.2 | 1.022 | 1.014 | 1.030 | 1.000 | 9535.2 | 11175.8 |
| XBB | 0.978 | 0.966 | 0.989 | 1.000 | 11931.6 | 12575.1 |
| BN.1.5 | 0.972 | 0.949 | 0.992 | 1.000 | 13327.3 | 10945.0 |
| XBB.3 | 0.954 | 0.938 | 0.969 | 1.000 | 12625.8 | 11771.7 |
| BM.1.1.3 | 0.883 | 0.837 | 0.924 | 1.000 | 14763.7 | 11523.7 |
| BA.2.75.2 | 0.875 | 0.829 | 0.918 | 1.000 | 15452.5 | 11566.5 |

The relative Re of XBB.1 is set to 1.

**Table S4. Primers used in this study**

| Primer name | Primer sequence (5'-to-3') | Purpose |
|---|---|---|
| Omicron universal Fw | cactatagggcgaattgggtaccatgtttgtgttcctggt | Preparation of S expression plasmid |
| BA2 Rv | agctccaccgcggtggcggccgctcaggtgtagtgcagtttca | Preparation of S expression plasmid |
| E180V Fw | ttcctgatggacttgGTGggcaagGAGggcaa | Preparation of S expression plasmid |
| E180V Rv | ttgccCTCcttgccCACcaagtccatcaggaa | Preparation of S expression plasmid |
| K478R Fw | accaggctggcaacAGGccatgtaatggagtg | Preparation of S expression plasmid |
| K478R_Rv | cactccattacatggCCTgttgccagcctggt | Preparation of S expression plasmid |

### Supplementary figure

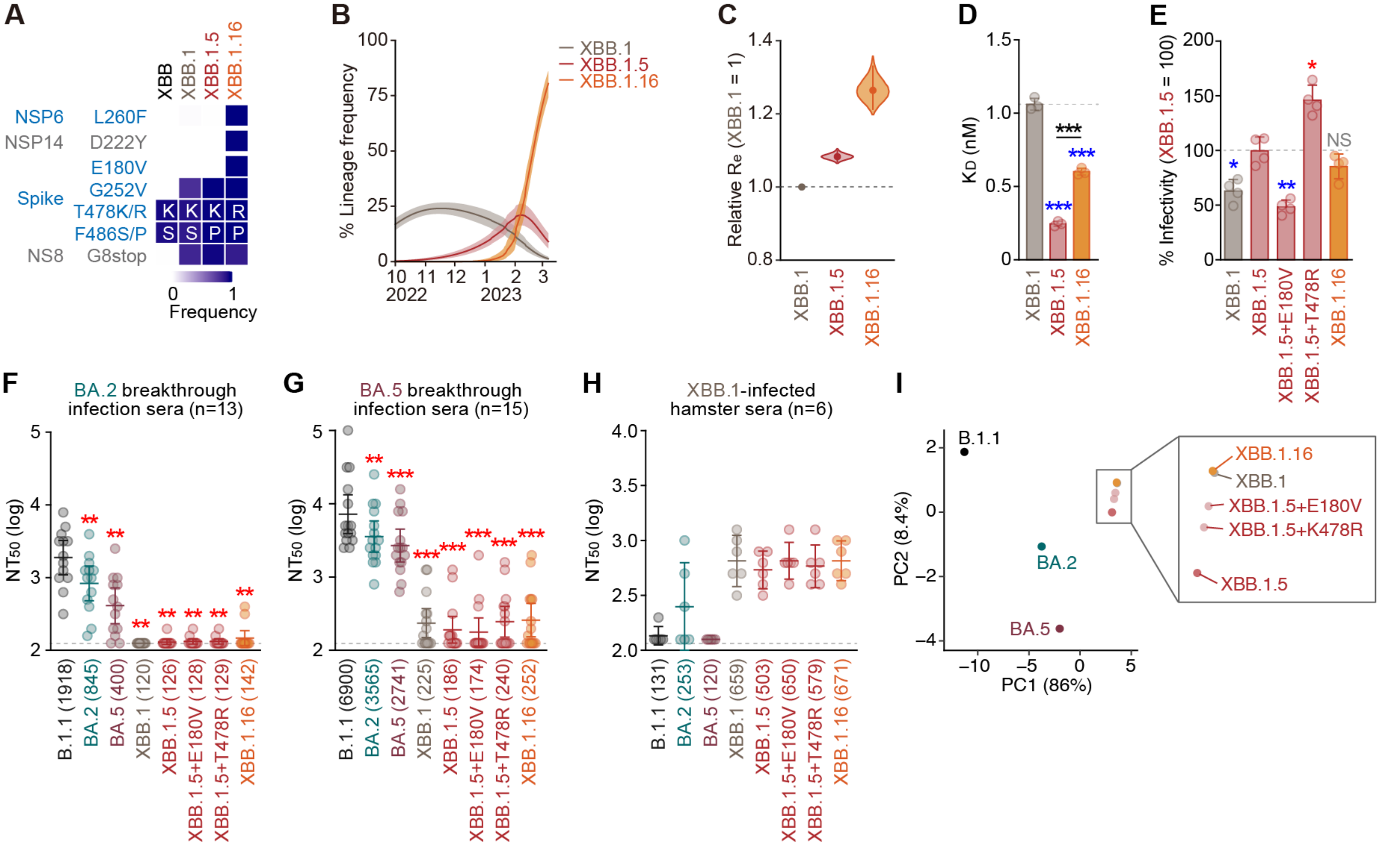


**Figure S1. Virological features of XBB.1.16**

(**A**) Frequency of mutations of interest in the representative XBB sublineages. Only mutations with a frequency >0.9 in at least one but not all the representative sublineages are shown.

(**B**) Estimated epidemic dynamics of the representative XBB sublineages in India from October 1, 2022 to March 5, 2023. Line, posterior mean; ribbon, 95% Bayesian confidence interval.

(**C**) Estimated relative $R_e$ of the representative XBB sublineages. The relative $R_e$ of XBB.1 is set to 1 (horizontal dashed line). Violin, posterior distribution; dot, posterior mean; line, 95% Bayesian confidence interval.

(**D**) Binding affinity of the RBD of SARS-CoV-2 S protein to the human ACE2 receptor by yeast surface display. The dissociation constant ($K_D$) value indicating the binding affinity of the RBD of the SARS-CoV-2 S protein to soluble ACE2 when expressed on yeast is shown. The horizontal dashed line indicates the mean $K_D$ value of the RBD of the XBB.1.

(**E**) Lentivirus-based pseudovirus assay. HOS-ACE2-TMPRSS2 cells were infected with pseudoviruses bearing each S protein. The amount of input virus was normalized to the amount of HIV-1 p24 capsid protein. The percentage infectivity of XBB.1, XBB.1.5+E180V (XBB.1+S:F486P/E180V), XBB.1.5+T478R (XBB.1+S:F486P/T478R), and XBB.1.16 (XBB.1+S:F486P/E180V/T478R) compared to that of XBB.1.5 are shown. The horizontal dashed line indicates the mean value of the percentage infectivity of the XBB.1.5. Assays were performed in quadruplicate. The presented data are expressed as the average ± SD. Each dot indicates the result of an individual replicate. Red and blue asterisks, respectively, indicate decreased and increased percentage infectivity. NS, no statistical significance.

(**F-H**) Neutralization assay. Assays were performed with pseudoviruses harboring the S proteins of B.1.1, BA.2, BA.5, XBB.1, XBB.1.5 (XBB.1+S:F486P), XBB.1.5+E180V (XBB.1+S:F486P/E180V), XBB.1.5+T478R (XBB.1+S:F486P/T478R), and XBB.1.16 (XBB.1+S:F486P/E180V/T478R). The following sera were used. (**F and G**) Convalescent sera from fully vaccinated individuals who had been infected with BA.2 after full vaccination (9 2-dose vaccinated and 4 3-dose vaccinated. 13 donors in total) (**F**) and those who had been infected with BA.5 after full vaccination (1 2-dose vaccinated donors, 13 3-dose vaccinated donors and 1 4-dose vaccinated donors. 15 donors in total) (**G**) and sera from hamsters infected with XBB.1 (6 hamsters)[9] (**H**) were used. Each dot indicates the result of an individual replicate. Assays for each serum sample were performed in triplicate to determine the 50% neutralization titer (NT50). Each dot represents one NT50 value, and the geometric mean and 95% confidence interval are shown. The number in parenthesis indicates the geometric mean of $NT_{50}$ values. The horizontal dashed line indicates the detection limit (120-fold).

(**I**) Principal component (PC) analysis representing the antigenicity of the S proteins. The analysis is based on the results of neutralization assays using BA.2 and BA.5 breakthrough infection sera and XBB.1-infected hamster's sera (**F–H**).

In **E**, statistically significant differences (*, P < 0.01, **, P < 0.001) versus XBB.1.5 were determined by two-sided Student's t tests. In **F and G**, statistically significant differences (**, P < 0.001, ***, P < 0.0001) versus B.1.1 were determined by two-sided Wilcoxon signed-rank tests and indicated with asterisks. Red asterisks indicate decreased NT50s. Information on the convalescent donors is summarized in **Table S2**.

### Consortia

#### The Genotype to Phenotype Japan (G2P-Japan) Consortium

**The Institute of Medical Science, The University of Tokyo, Japan**

Yu Kaku, Naoko Misawa, Ziyi Guo, Alfredo A. Hinay Jr., Shigeru Fujita, Jarel Elgin M. Tolentino, Luo Chen, Mai Suganami, Mika Chiba, Ryo Yoshimura, Kyoko Yasuda, Keiko Iida, Naomi Ohsumi, Adam P. Strange

**Hokkaido University, Japan**

Takasuke Fukuhara, Tomokazu Tamura, Rigel Suzuki, Saori Suzuki, Hayato Ito, Keita Matsuno, Hirofumi Sawa, Naganori Nao, Shinya Tanaka, Masumi Tsuda, Lei Wang, Yoshikata Oda, Zannatul Ferdous, Kenji Shishido

**Tokyo Metropolitan Institute of Public Health, Japan**

Kenji Sadamasu, Kazuhisa Yoshimura, Hiroyuki Asakura, Isao Yoshida, Mami Nagashima

**Tokai University, Japan**

So Nakagawa

**Kyoto University, Japan**

Kotaro Shirakawa, Akifumi Takaori-Kondo, Kayoko Nagata, Ryosuke Nomura, Yoshihito Horisawa, Yusuke Tashiro, Yugo Kawai, Kazuo Takayama, Rina Hashimoto, Sayaka Deguchi, Yukio Watanabe, Ayaka Sakamoto, Naoko Yasuhara, Takao Hashiguchi, Tateki Suzuki, Kanako Kimura, Jiei Sasaki, Yukari Nakajima, Hisano Yajima

**Hiroshima University, Japan**

Takashi Irie, Ryoko Kawabata

**Kyushu University, Japan**

Kaori Tabata

**Kumamoto University, Japan**

Terumasa Ikeda, Hesham Nasser, Ryo Shimizu, MST Monira Begum, Michael Jonathan, Yuka Mugita, Otowa Takahashi, Kimiko Ichihara, Takamasa Ueno, Chihiro Motozono, Mako Toyoda

**University of Miyazaki, Japan**

Akatsuki Saito, Maya Shofa, Yuki Shibatani, Tomoko Nishiuchi

**Acknowledgments**

We would like to thank all members of The Genotype to Phenotype Japan (G2P-Japan) Consortium. We thank Dr. Kenzo Tokunaga (National Institute of Infectious Diseases, Japan) for sharing materials and Dr. Jin Kuramochi (Interpark Kuramochi Clinic, Japan) for sharing human sera. We gratefully acknowledge the numerous laboratories worldwide that have provided sequence data and metadata to GISAID. A full list of originating and submitting laboratories for the sequences used in our analysis can be found at https://www.gisaid.org using the EPI-SET-ID: EPI_SET_230403zh.

### Supplementary References


1. Yamasoba D, Kimura I, Nasser H, et al. Virological characteristics of the SARS-CoV-2 Omicron BA.2 spike. *Cell* 2022; **185**(12): 2103-15.e19.
2. Kimura I, Yamasoba D, Tamura T, et al. Virological characteristics of the novel SARS-CoV-2 Omicron variants including BA.4 and BA.5. *Cell* 2022; **185**(21): 3992-4007.e16.
3. Saito A, Tamura T, Zahradnik J, et al. Virological characteristics of the SARS-CoV-2 Omicron BA.2.75 variant. *Cell Host Microbe* 2022; **30**(11): 1540–55.e15.
4. Gu Z, Eils R, Schlesner M. Complex heatmaps reveal patterns and correlations in multidimensional genomic data. *Bioinformatics* 2016; **32**(18): 2847-9.
5. Motozono C, Toyoda M, Zahradnik J, et al. SARS-CoV-2 spike L452R variant evades cellular immunity and increases infectivity. *Cell Host Microbe* 2021; **29**(7): 1124-36.
6. Uriu K, Kimura I, Shirakawa K, et al. Neutralization of the SARS-CoV-2 Mu variant by convalescent and vaccine serum. *N Engl J Med* 2021; **385**(25): 2397-9.
7. Yamasoba D, Kosugi Y, Kimura I, et al. Neutralisation sensitivity of SARS-CoV-2 omicron subvariants to therapeutic monoclonal antibodies. *Lancet Infect Dis* 2022; **22**(7): 942-3.
8. Ito J, Suzuki R, Uriu K, et al. Convergent evolution of the SARS-CoV-2 Omicron subvariants leading to the emergence of BQ.1.1 variant. *BioRxiv* 2022: doi: https://doi.org/10.1101/2022.12.05.519085.
9. Tamura T, Ito J, Uriu K, et al. Virological characteristics of the SARS-CoV-2 XBB variant derived from recombination of two Omicron subvariants. *BioRxiv* 2022: doi: https://doi.org/10.1101/2022.12.27.521986.
10. Uriu K, Ito J, Zahradnik J, et al. Enhanced transmissibility, infectivity, and immune resistance of the SARS-CoV-2 omicron XBB.1.5 variant. *Lancet Infect Dis* 2023; **23**(3): 280-1.
11. Niwa H, Yamamura K, Miyazaki J. Efficient selection for high-expression transfectants with a novel eukaryotic vector. *Gene* 1991; **108**(2): 193-9.
12. Zahradnik J, Dey D, Marciano S, et al. A protein-engineered, enhanced yeast display platform for rapid evolution of challenging targets. *ACS Synth Biol* 2021; **10**(12): 3445-60.
13. Zahradnik J, Marciano S, Shemesh M, et al. SARS-CoV-2 variant prediction and antiviral drug design are enabled by RBD in vitro evolution. *Nat Microbiol* 2021; **6**(9): 1188-98.
14. Kimura I, Kosugi Y, Wu J, et al. The SARS-CoV-2 Lambda variant exhibits enhanced infectivity and immune resistance. *Cell Rep* 2022; **38**(2): 110218.
15. Ferreira I, Kemp SA, Datir R, et al. SARS-CoV-2 B.1.617 mutations L452R and E484Q are not synergistic for antibody evasion. *J Infect Dis* 2021; **224**(6): 989-94.
16. Ozono S, Zhang Y, Ode H, et al. SARS-CoV-2 D614G spike mutation increases entry efficiency with enhanced ACE2-binding affinity. *Nat Commun* 2021; **12**(1): 848.
17. Fujita S, Kosugi Y, Kimura I, Yamasoba D, The Genotype to Phenotype Japan (G2P-Japan) Consortium, Sato K. Structural Insight into the Resistance of the SARS-CoV-2 Omicron BA.4 and BA.5 Variants to Cilgavimab. *Viruses* 2022; **14**(12): 2677.

18. Uriu K, Cardenas P, Munoz E, et al. Characterization of the immune resistance of SARS-CoV-2 Mu variant and the robust immunity induced by Mu infection. *J Infect Dis* 2022.
19. Saito A, Irie T, Suzuki R, et al. Enhanced fusogenicity and pathogenicity of SARS-CoV-2 Delta P681R mutation. *Nature* 2022; **602**(7896): 300-6.
20. Kimura I, Yamasoba D, Nasser H, et al. The SARS-CoV-2 spike S375F mutation characterizes the Omicron BA.1 variant. *iScience* 2022; **25**(12): 105720.
21. Ozono S, Zhang Y, Tobiume M, Kishigami S, Tokunaga K. Super-rapid quantitation of the production of HIV-1 harboring a luminescent peptide tag. *J Biol Chem* 2020; **295**(37): 13023-30.
